# Aggrecan hypomorphism accelerates the progression of post-traumatic osteoarthritis in mice

**DOI:** 10.64898/2026.08.06.742527

**Authors:** Xujia Wang, Reinhild Hofmann, Bastian Hartmann, Shenxi Zhong, Xiangqing Meng, Heidrun Grondinger, Zsuzsanna Farkas, Riccardo E Giunta, Hauke Clausen-Schaumann, Attila Aszodi, Paolo Alberton

**Affiliations:** Musculoskeletal University Center Munich (MUM), Department of Orthopaedics and Trauma Surgery, Ludwig-Maximilians-University (LMU), Munich, Germany; Center for Applied Tissue Engineering and Regenerative Medicine, Munich University of Applied Sciences, Munich, Germany; Division of Hand, Plastic and Aesthetic Surgery, LMU University Hospital, LMU Munich, Germany

**Keywords:** aggrecan, extracellular matrix, post-traumatic osteoarthritis, articular cartilage, cartilage biomechanics

## Abstract

**Objective:** The *Agc1^CreERT2^* mouse line is a powerful tamoxifen-inducible genetic tool used for conditional gene manipulation specifically in cartilage. The aim of this study was to investigate the effects of aggrecan hypomorphism on the progression of post-traumatic osteoarthritis (PT-OA) in *Agc1^CreERT2^* mice.

**Methods:** Proteoglycan content in cartilage samples from the knees of E18.5 embryos were quantified by sulfated glycosaminoglycan (sGAG) assay. Destabilization of the medial meniscus (DMM) surgery was performed to induce PT-OA in 12-week-old wild-type, heterozygous *Agc1^CreERT2/+^* and homozygous *Agc1^CreERT2/CreERT2^* mice. Progression of OA was assessed at 4-, 8-, and 12-weeks post-DMM by OARSI, synovitis and osteophyte maturation histopathology scores and micro-computed tomography (µCT). Aggrecan deposition, cartilage matrix-degrading proteases, aggrecan and collagen II degradation neoepitopes were investigated by immunohistochemical staining, and serum C-terminal cross-linked telopeptide of type II collagen (CTX-II) levels by an enzyme-linked immunosorbent assay (ELISA). Chondrocyte apoptosis was analyzed with the terminal deoxynucleotidyl transferase (TdT) dUTP nick-end labeling (TUNEL) assay. The biomechanical properties of articular cartilage (AC) were investigated with indentation-type atomic force microscopy (IT-AFM).

**Results:** Before DMM, homozygous *Agc1^CreERT2/CreERT2^* mice had reduced sGAG and aggrecan levels and increased cartilage stiffness. After DMM, they exhibited increased cartilage degradation, synovitis, osteophyte formation and meniscus mineralization, chondrocyte apoptosis and cartilage stiffness compared with wild-type mice. Immunohistochemistry demonstrated increased expression of the aggrecanase ADAMTS-5, the metalloproteinase MMP-13, the aggrecan degradation neoepitope NITEGE and the collagen degradation neoepitope C1,2C in AC. ELISA also revealed elevated serum CTX-II levels. Heterozygous *Agc1^CreERT2/+^* mice also exhibited accelerated PT-OA compared with wild-type mice, characterized by elevated CTX-II levels at 4-weeks, increased synovitis, osteophyte and soft tissue mineralization at 8-weeks, and more severe cartilage degeneration at 12-weeks post-DMM.

**Conclusion:** Both homozygous and heterozygous *Agc1^CreERT2^*mice exhibit increased susceptibility to PT-OA, underscoring the importance of physiological aggrecan expression in maintaining joint homeostasis and regulating joint pathophysiology. These findings indicate that *Agc1^CreERT2/+^*mice are not phenotypically neutral in the DMM model and that this intrinsic susceptibility should be considered when interpreting studies employing inducible, cartilage-specific gene deletion.

## Introduction

Post-traumatic osteoarthritis (PT-OA) is a debilitating condition that typically develops following joint trauma such as sports injuries, accidents, or surgical interventions (1). Unlike idiopathic osteoarthritis (OA), which usually develops as a result of aging, PT-OA can affect younger people and progresses more rapidly. This condition not only affects quality of life due to pain and disability, but also imposes significant healthcare costs (2,3).

Aggrecan, the major proteoglycan of the cartilage extracellular matrix (ECM), is essential for cartilage biomechanics. Its highly negatively charged glycosaminoglycan (GAG) chains retain water, enabling the tissue to resist compressive loading. Aggrecan loss compromises ECM integrity, reducing its shock-absorbing capacity and compromising joint stability, key factors in the pathogenesis of both OA and PT-OA (4). Following joint injury, increased aggrecanases (ADAMTS-4 and −5) and matrix metalloproteinase (MMP) activity rapidly cleave the aggrecan core protein, causing GAG loss, exposing the collagen network to damage, and disrupting cellular mechanotransduction, thereby accelerating cartilage degeneration (5).

Mutations in the human gene encoding aggrecan (*ACAN*) lead to aggrecanopathies, including spondyloepiphyseal dysplasia with short stature and severe OA, autosomal dominant familial osteochondritis dissecans with early OA, and various idiopathic short stature syndromes associated with accelerated bone maturation (6–9). *ACAN* mutations are mostly dominant negative missense mutations that disrupt the normal function of the protein, or frameshift mutations that result in premature termination codons and aggrecan protein haploinsufficiency (7–9). A recent large-scale genome-wide association study meta-analysis identified a missense *ACAN* variant associated with knee and hip OA, suggesting that ACAN plays a role in common idiopathic OA (10).

The *Agc1^tm(IRES-CreERT2)^* mouse line (*Agc1^CreERT2^*) was generated for conditional inactivation of floxed genes in chondrocytes by inserting a tamoxifen-inducible *CreERT2* cassette into the 3’ untranslated region (UTR) of the mouse aggrecan gene (*Agc1*). This knock-in strategy resulted in a hypomorphic, partial-loss-of-function mutation due to a 50%-64% reduction in *Agc1* mRNA levels in chondrocytes and about a 50% reduction of aggrecan protein in the cartilage of homozygous *Agc1^CreERT2/CreERT2^*mice (11,12). *Agc1^CreERT2/CreERT2^* mice displayed postnatal dwarfism associated with a shortened growth plate and AC (11,12). Furthermore, we have shown that the reduced aggrecan levels in *Agc1^CreERT2/CreERT2^* mice lead to increased nano-stiffness of the AC ECM and to severe spontaneous OA at 12 months of age (12). Of note, heterozygous *Agc1^CreERT2/+^* mice also exhibited mild, age- and sex-dependent growth abnormalities.

In this study, we employed the DMM model (13) to induce PT-OA in *Agc1^CreERT2/CreERT2^* mice and determine how reduced aggrecan levels influence PT-OA progression.

## Methods

### Experimental workflow

**Figure 1.**
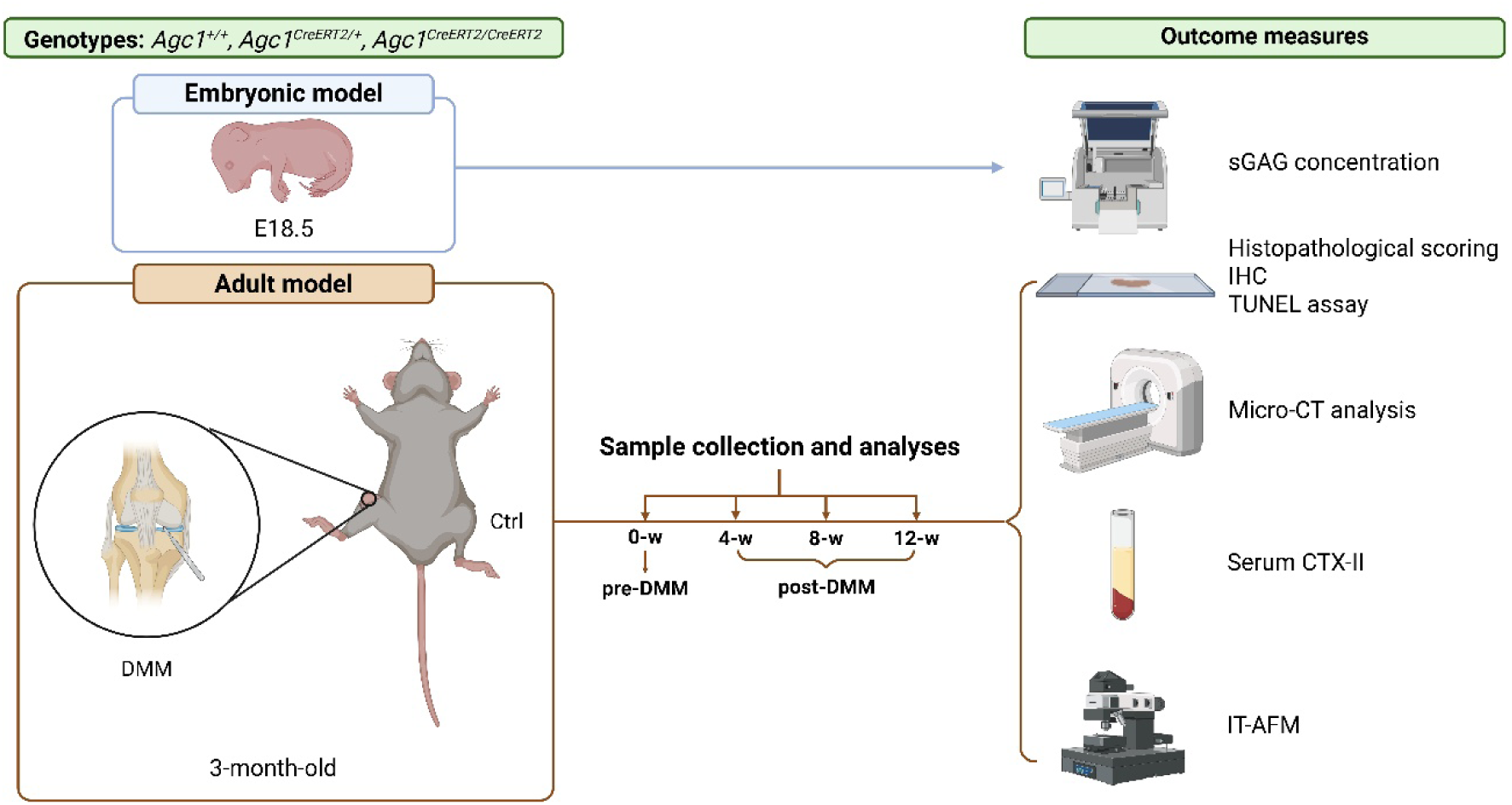
Overview of the experimental design and outcome measures. Three genotypes were included: *Agc1^+/+^, Agc1^CreERT2/+^, and Agc1^CreERT2/CreERT2^*. Upper panel: Embryonic assessment. Knee cartilage was collected from embryos at embryonic day 18.5 (E18.5), and sGAG content was quantified using the Blyscan assay and normalized to cartilage wet weight. Lower panel: Adult DMM model. 3-month-old male mice were assigned to a pre-DMM group sampled at 0-week(-w) or underwent DMM surgery in the right knee, with the contralateral left knee serving as the control. Samples were collected before DMM surgery at 0-w and at 4-, 8-, and 12-w after surgery. Outcome measures. Joint pathology and cartilage degeneration were evaluated by histopathological scoring, IHC, TUNEL, µCT, CTX-II measurement, and IT-AFM. Detailed experimental details are provided in the Methods section. Figure created with BioRender.com.

### Experimental animals

The *Agc1^CreERT2^* mouse line (14) was obtained from the Jackson Laboratory (JAX stock #019148). Animals were maintained in plastic cages in an air- and temperature-controlled environment under an alternating 12:12-h light-dark cycle. All mouse experiments were performed according to the federal institutional guidelines for the care and use of laboratory animals, approved by the Central Animal Facility of LMU Munich and the Government of Upper Bavaria (Application number: ROB-55.2-2532.Vet_02-17-155).

### DMM surgery

DMM operations were performed on the right knee joint of 12-week-old male mice. Mice were anaesthetized through intraperitoneal injection of an anesthetic mixture (medetomidine, 0.5 mg/kg; midazolam, 5.0 mg/kg; fentanyl, 0.05 mg/kg). The surgery was performed as described in Glasson et al. (13). Contralateral left knee joints were used as the control group. Mice were sacrificed by intraperitoneal injection overdose of a xylazine (30 mg/kg) and ketamine (150 mg/kg) mixture, at the time points of 4-, 8- and 12-w post-operation.

### Micro-computed tomography (µCT) analysis

Deskinned hind limbs of euthanized mice were scanned in air using a ProCon X-Ray µCT device (CT-ALPHA, ProCon X-Ray GmbH, Sarstedt, Germany; DFG number: INST 409_211-1) with a voxel size of 6 µm (100 kV, 200 µA, 0.5 mm aluminum filter, 0.24° rotation angle). The projection images were corrected for imaging artifacts using X-AID post-processing software (version 2022.7.0; MITOS GmbH, Munich, Germany).

For trabecular bone analysis, a volume of interest (VOI) was defined in the medial tibial subchondral bone to exclude the subchondral bone plate, cortical bone, and growth plate (15). Bone volume fraction (BV/TV), trabecular thickness (Tb.Th), and trabecular separation (Tb.Sp) were quantified within this region. Ectopic mineralization was assessed by quantifying osteophytes, calcified menisci, and calcified periarticular soft tissues. All measurements were performed using Dragonfly 3D visualization and image analysis software (version 2022.1.0.1249; Object Research Systems Inc., Montreal, Canada).

### Sample preparation, histopathology and immunohistochemistry

DMM and contralateral control hind limbs were dissected, skinned, fixed overnight in 4% paraformaldehyde in phosphate-buffered saline (PBS) at 4°C, decalcified in 20% EDTA (pH 8.0; Sigma-Aldrich, St. Louis, MO, USA), and paraffin-embedded. Coronal sections (8 µm) were prepared using an HM355S STS microtome (Thermo Scientific, Madison, WI, USA) and mounted on Superfrost Plus slides (Thermo Scientific). Every tenth section was stained with Safranin O/Fast Green or Toluidine Blue to evaluate the entire knee joint.

Cartilage degeneration in the medial tibial plateau (MTP) and the medial femoral condyle (MFC) was graded using the Osteoarthritis Research Society International (OARSI) histopathological scoring system (16). Synovitis was assessed using a modified published grading system (17), scoring synovial hyperplasia (graded 0-3), cellularity (graded 0-3) and fibrosis (graded 0-1) in the MTP and MFC; summed scores were used to determine overall synovitis severity. Osteophyte maturation was graded from 0 (none) to 5 (predominantly bone) as previously described (18). All histopathological assessments were performed independently by three blinded observers.

Immunohistochemistry was performed on deparaffinized and rehydrated sections using the avidin–biotin complex (ABC) method as previously described (12). Primary antibodies included rabbit anti-aggrecan (MilliporeSigma, AB1031, 1:200; Burlington, MA, USA), mouse anti-collagen type II (DSHB CIIC1, 5 µg/mL), rabbit anti-NITEGE (Novus Biologicals, NB100-74350, 1:500; Centennial, CO, USA), mouse anti-MMP-13 (Sigma-Aldrich, MAB13424, 1:100; St. Louis, MO, USA), rabbit anti-ADAMTS-5 (Abcam, ab13976, 1:500; Cambridge, UK), rabbit anti-VDIPEN (a gift from Amanda Fosang, University of Melbourne, 1:1000), and C1,2C (IBEX Technologies Inc., 50-1035, 1:400; Montreal, QC, Canada).

### Terminal deoxynucleotidyl transferase dUTP nick-end labeling (TUNEL) assay

Apoptotic cells were detected by TUNEL staining. Deparaffinized and rehydrated sections were permeabilized with 0.1% Triton X-100 and 0.1% sodium citrate for 8 min, rinsed with PBS, and air-dried. TUNEL staining was performed using the Fluorescein In Situ Cell Death Detection Kit (Roche, Mannheim, Germany) according to the manufacturer’s instructions. Nuclei were counterstained with DAPI-containing mounting medium (Fluoroshield; Abcam, Cambridge, UK). Images were acquired using a fluorescence microscope (Axio Observer; Carl Zeiss, Jena, Germany). TUNEL-positive (green) and DAPI-positive (blue) cells were quantified in the uncalcified and calcified zones of the AC using ImageJ software (National Institutes of Health, Bethesda, MD, USA). Apoptosis was expressed as the percentage of TUNEL-positive cells relative to the total number of DAPI-positive cells.

### Quantitative analysis of glycosaminoglycans

Cartilage samples from the knees of E18.5 embryos were weighed and digested with 125 µg/mL papain (Sigma-Aldrich) in papain buffer (0.1 M sodium acetate, pH 5.5, 5 mM EDTA, and 5 mM L-cysteine-HCl) for 5 h at 60°C. Following centrifugation at 10,000 × g for 5 min, sulfated glycosaminoglycan (sGAG) content in the supernatant was quantified using the Blyscan™ Sulfated Glycosaminoglycan Assay Kit (Biocolor Ltd., Carrickfergus, UK) according to the manufacturer’s instructions. Absorbance was measured at 656 nm using a microplate reader (Thermo Fisher Scientific), and sGAG concentrations were determined from a chondroitin 4-sulfate standard curve and normalized to the weight of the digested cartilage.

### C-telopeptide of type II collagen (CTX-II) assay

Blood samples (ca. 200 μl) were collected post-mortem from the inferior vena cava at 4-, 8-, and 12-w post-DMM. Samples were allowed to clot for 30 min at RT, centrifuged at 2,500 rpm for 15 min, and the serum was collected and stored at −80°C. Serum levels of the type II collagen degradation biomarker C-telopeptide of type II collagen (CTX-II) were quantified using a mouse CTX-II ELISA kit (MyBioSource MBS706197, San Diego, CA, USA) according to the manufacturer’s instructions.

### Indentation-type atomic force microscopy (IT-AFM)

Hind limbs were dissected immediately after euthanasia, skinned, embedded in Tissue-Tek O.C.T. compound (Sakura, Zoeterwoude, The Netherlands) without prior fixation, and gradually frozen on a dry ice-cooled copper plate. Cryosections (30 µm) of the knee joint were prepared using a Leica CM1950 cryostat and stored at −20°C until analysis. Nano-scale mechanical properties were assessed using indentation-type atomic force microscopy (IT-AFM) as previously described (12). Three force maps (3 × 3 µm; 25 × 25 force curves each) were acquired from the superficial, middle, and deep zones of the medial tibial plateau using a tip velocity of 15 µm/s and a maximum loading force of 12 nN. A total of 1,875 force curves per cartilage zone were recorded from each section, with two sections analyzed per animal. The Young’s modulus was derived from the force-indentation curves up to a maximum indentation depth of 1 µm, applying a modified Hertz-Sneddon model for a four-sided pyramidal indenter (19) using the CANTER Processing Toolbox (v. 5.6.1, GitHub, 2022, California, US. https://github.com/CANTERhm/CANTER_Processing_Tool). Histograms of Young’s modulus were generated for each post-DMM stage and genotype, and distribution peaks were determined by bimodal Gaussian fitting.

### Statistical analysis

Statistical analyses were carried out using GraphPad Prism-v10 software (GraphPad, San Diego, CA, USA). Statistical significance was calculated after determination of the Gaussian distribution and homogeneity of variance test using the Kruskal-Wallis test, one-way or two-way ANOVA tests (respectively for one or two independent variables), or Welch’s ANOVA test with appropriate post hoc tests. Between-group results are presented as mean differences (MD), 95% confidence interval (95% CI) and adjusted *p* values (*P*) ≤ 0.05 were considered statistically significant.

## Results

### Effects of reduced aggrecan levels on joint tissues before surgery

Prior to DMM, 3-month-old *Agc1^CreERT2/CreERT2^* mice exhibited reduced toluidine blue staining and aggrecan deposition in the cartilage ECM compared with *Agc1^CreERT2/+^* and *Agc1^+/+^*mice [Fig. 2(A, C)], accompanied by slightly thinner uncalcified and calcified AC thickness [Fig. 2(B)]. At embryonic day 18.5, cartilage sGAG was significantly lower in *Agc1^CreERT2/CreERT2^*than *Agc1^+/+^* (MD = −4.407 µg/mg, 95% CI: [-8.754, −0.05940], *P* = 0.0482) and *Agc1^CreERT2/+^* mice (MD = −5.826 µg/mg, 95% CI: [-10.89, −0.7637], *P* = 0.0316) [Fig. 2 (D)]. Baseline OARSI and osteophyte maturity scores were similar across genotypes, although osteophyte maturity showed a stepwise increase from wild-type to heterozygous and homozygous mice [Fig. 2(E, F)]. Synovitis was mildly but significantly increased in *Agc1^CreERT2/CreERT2^* mice compared with wild-type mice (mean rank difference = 6.5, *P* = 0.0190) [Fig. 2(G, H)], whereas chondrocyte apoptosis did not differ between genotypes [Fig. 2(I, J)]. To assess the biomechanical consequences of reduced proteoglycan content, AC was analyzed by nanoscale IT-AFM [Fig. 2(K)]. *Agc1^CreERT2/CreERT2^* cartilage displayed markedly increased superficial-zone stiffness compared with wild type, with 3.4- and 4.1-fold increases in the proteoglycan-associated modulus (E1) and collagen-network-associated modulus (E2), respectively. Notably, *Agc1^CreERT2/CreERT2^*cartilage lacked the normal depth-dependent increase in stiffness, with E1 and E2 remaining comparable across all cartilage zones. The superficial zone of *Agc1^CreERT2/CreERT2^* cartilage also displayed a substantially broader stiffness distribution (0.03–5.99 MPa) than wild-type cartilage (0.03–2.57 MPa), whereas heterozygous cartilage showed only a moderate broadening (0.04–3.47 MPa). Aside from this greater superficial-zone variability, Young’s modulus distributions were comparable between *Agc1^+/+^* and *Agc1^CreERT2/+^* across all cartilage zones.

**Figure 2.**
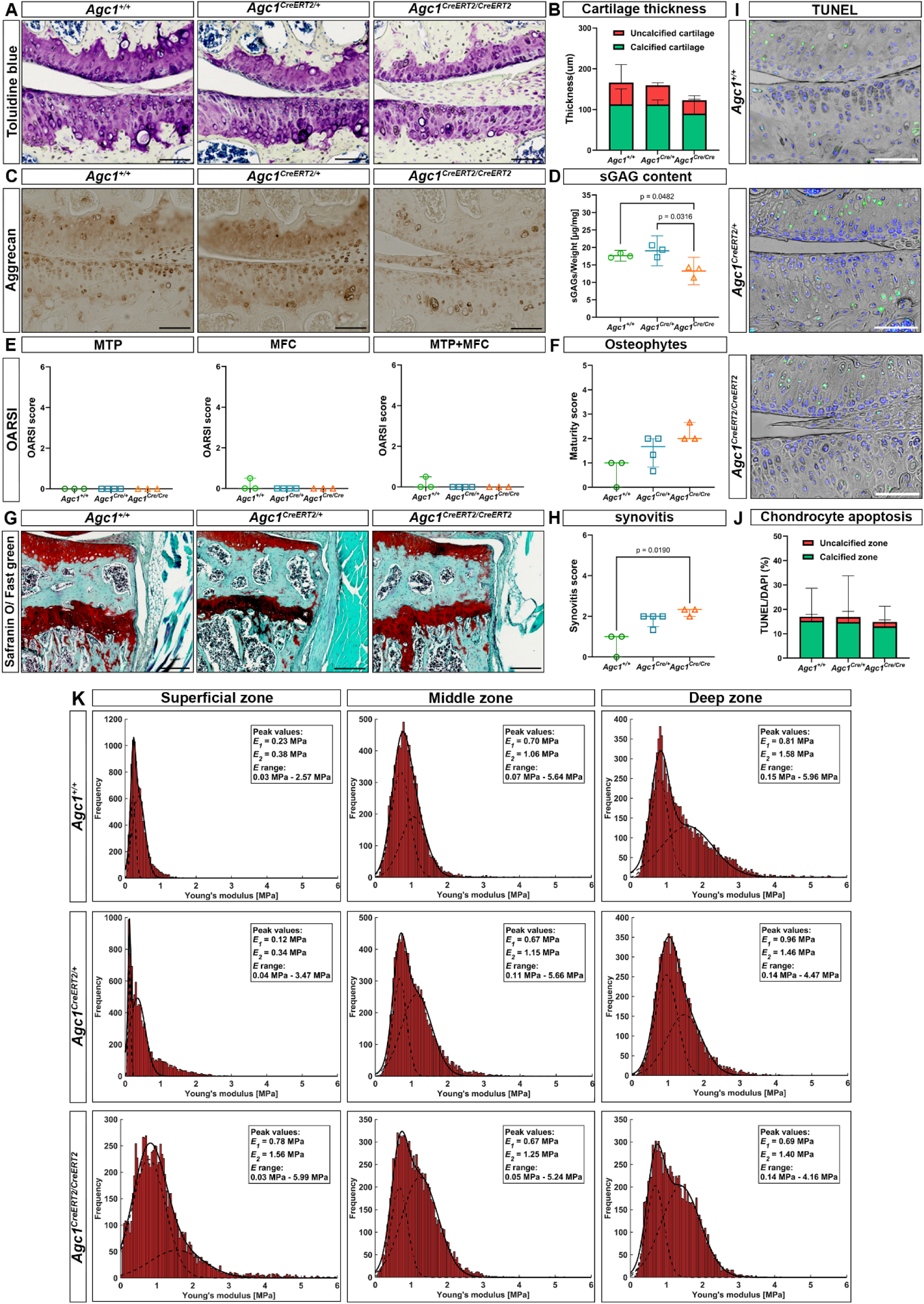
Baseline histological and biochemical assessment of knee joints in 3-month-old mice. (A) Representative toluidine blue stained sections of medial tibiofemoral articular cartilage from *Agc1^+/+^*, *Agc1^CreERT2/+^*, and *Agc1^CreERT2/CreERT2^*mice. Scale bars = 50 μm. (B) Quantification of articular cartilage thickness, partitioned into uncalcified and calcified cartilage layers. (C) Representative aggrecan immunohistochemistry in articular cartilage. Scale bars=50 μm. (D) sGAG content normalized to cartilage weight, in E18,5 developing knee joints. (E) OARSI scores for the AC of the MTP, MFC, and combined MTP+MFC regions. (F) Modified osteophyte maturity score in the knee joint. (G) Representative Safranin O/Fast Green-stained knee sections used for synovitis scoring. Scale bars=200 μm, and (H) synovitis score. (I) Representative TUNEL staining (green) with DAPI nuclear counterstain (blue). Scale bars = 50 μm. (J) Quantification of TUNEL positive chondrocytes, analyzed separately in uncalcified and calcified cartilage zones. In Fig. 2(B, E, F, H and J), data are shown as median with interquartile range, statistical comparisons among genotypes were performed using the Kruskal–Wallis test followed by Dunn’s multiple-comparisons post hoc test (n ≥ 3); in Fig. 2D, the graph data are shown as mean ± 95% CI. Groups were compared using Welch’s one-way ANOVA followed by Dunnett’s T3 multiple-comparisons test (n = 3). Exact *p* values are shown where indicated. (K) Analysis of the biomechanical properties of the AC in 3-month-old Agc1^+/+^, *Agc1^CreERT2/+^* and *Agc1^CreERT2/CreERT2^*mice. Young’s Modulus histograms illustrating the stiffness distribution in the three AC zones of the tibial plateau determined by IT-AFM (n = 3). In each histogram, the solid line represents the combination of two Gaussian functions, while the dashed lines indicate individual fits, namely E1 and E2, corresponding to the proteoglycan gel and the collagen fibrils, respectively. Note the drastically increased E1 and E2 values in the superficial zone of *Agc1^CreERT2/CreERT2^*mice compared to *Agc1^+/+^* and *Agc1^CreERT2/+^* animals.

### *Agc1^CreERT2/CreERT2^* mice develop severe AC degeneration after DMM

To investigate the impact of reduced aggrecan levels on PT-OA progression, 3-month-old male mice underwent DMM surgery, and knee joint pathology was assessed at 4-, 8-, and 12-w post-operatively [Fig. 3(A, B)]. At 4-w post-DMM, *Agc1^CreERT2/CreERT2^* knees exhibited significantly higher MTP OARSI scores than wild-type and heterozygous, with MDs of 2.088 (95% CI: [0.8493, 3.326], *P* = 0.0006) and 1.687 (95% CI: [0.6321, 2.743], *P* = 0.0011), respectively. In the MFC, homozygous knees also scored higher than heterozygous (MD = 1.178, 95% CI: [0.1127, 2.244], *P* = 0.0274). Combined MTP and MFC scores were significantly increased in *Agc1^CreERT2/CreERT2^* mice, compared with wild-type (MD = 3.325, 95% CI: [1.038, 5.612], *P* = 0.0029), *Agc1^CreERT2/+^* (MD = 2.866, 95% CI: [0.9161, 4.815], *P* = 0.0026) and contralateral controls (MD = 4.144, 95% CI: [2.525, 5.762], *P* < 0.0001).

**Figure 3.**
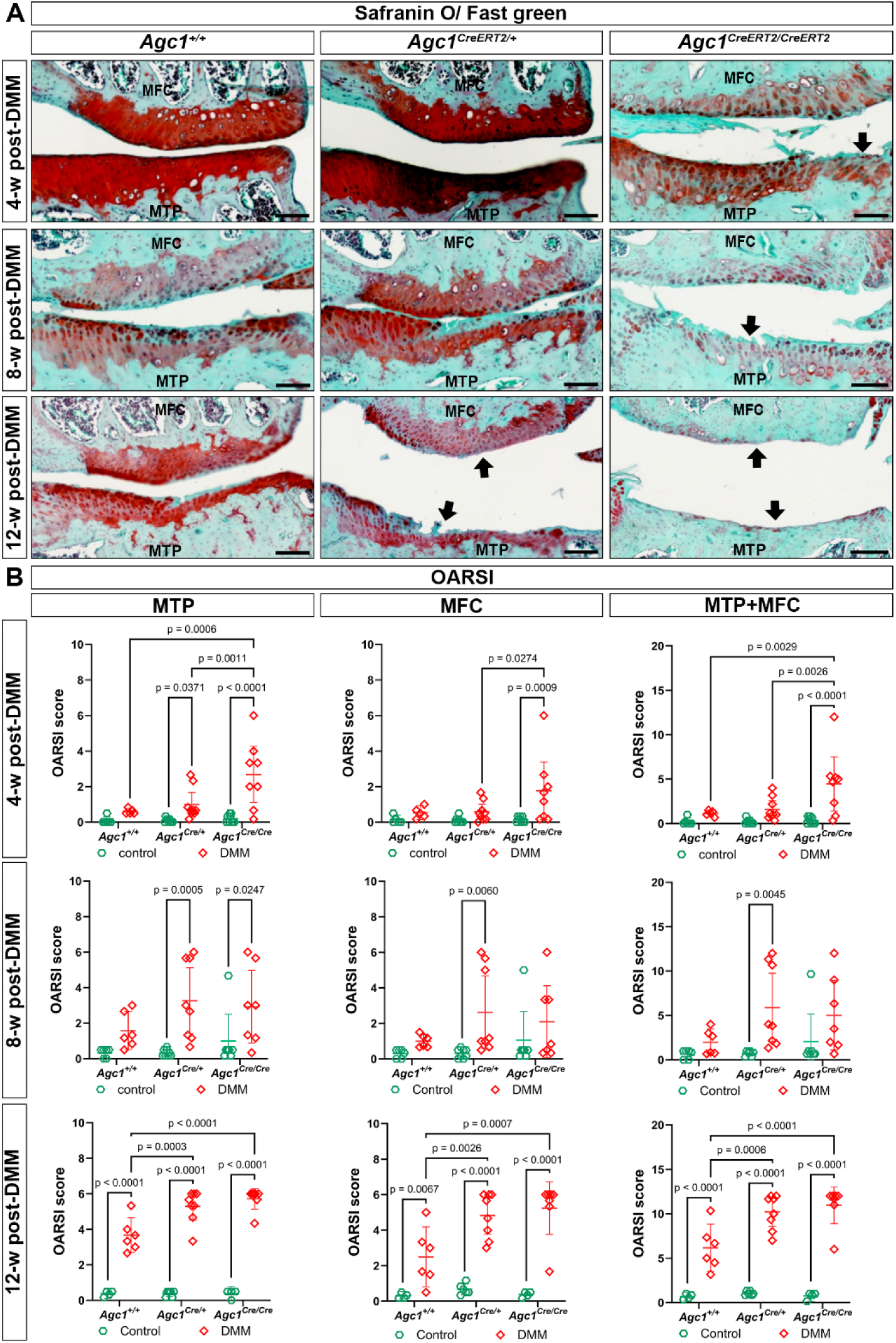
Articular cartilage degeneration following DMM surgery assessed by Safranin O/Fast Green staining and OARSI scoring. (A) Safranin O/Fast Green– stained knee joint sections showing varying degrees of cartilage degeneration at 4-, 8-, and 12-w post–DMM in *Agc1^+/+^*, *Agc1^CreERT2/+^*, and *Agc1^CreERT2/CreERT2^*mice. Arrows indicate regions of cartilage erosion. (B) Knee cartilage degeneration quantified by the OARSI score for the MTP, MFC, and combined MTP+MFC at 4-, 8-, and 12-w post-DMM; contralateral knees served as controls. The graph data are shown as mean ± 95% CI. Statistical comparisons were performed using two-way ANOVA followed by Tukey’s multiple-comparisons post hoc test (n ≥ 4). Exact p values are shown in the plots. Scale bar = 200 μm.

At 8-w post-DMM, both *Agc1^CreERT2/+^* and *Agc1^CreERT2/CreERT2^*mice exhibited higher OARSI scores in the MTP and MFC compared to the wild-type mice; however, these differences did not reach statistical significance. Significant DMM-associated increases relative to the corresponding contralateral joints remained evident in the MTP of both mutant genotypes, whereas significant MFC and combined MTP+MFC changes were mainly detected in heterozygous mice.

By 12-w, all DMM joints displayed advanced medial compartment degeneration with significantly higher OARSI scores than the corresponding contralateral joints. Importantly, *Agc1^CreERT2/CreERT2^* mice exhibited significantly more severe cartilage erosion than wild-type mice in both MTP (MD = 2.048, 95% CI: [1.126, 2.969], *P* < 0.0001) and MFC (MD = 2.738, 95% CI: [1.127,4.349], *P* = 0.0007). Interestingly, heterozygous mice also displayed significantly higher OARSI scores than wild-type after DMM in both MTP (MD = 1.625, 95% CI: [0.7303, 2.520], *P* = 0.0003) and MFC (MD = 2.333, 95% CI: [0.7695, 3.897], p = 0.0026). Moreover, combined MTP+MFC OARSI scores confirmed a genotype-dependent aggravation of PT-OA, with higher scores in *Agc1^CreERT2/+^* mice (MD = 4.042, 95% CI: [1.702, 6.381], *P* = 0.0006) and the greatest degeneration in *Agc1^CreERT2/CreERT2^*mice (MD = 4.786, 95% CI: [2.376, 7.195], *P* < 0.0001) relative to wild-type mice. Contralateral joints remained largely unaffected.

### *Agc1^CreERT2/CreERT2^* mice exhibit enhanced DMM-induced synovitis and accelerated osteophyte maturation

We next evaluated synovitis and osteophyte maturation as additional pathological features of OA [Fig. 4(A-D)]. *Agc1^CreERT2/CreERT2^*mice showed significantly increased post-DMM synovitis than wild-type mice at 4- and 8-w, with MDs of 2.083 (95% CI: [0.8099, 3.357], *P* = 0.0008) and 2.429 (95% CI: [0.8579, 3.999], *P* = 0.0016). At 8-w post-DMM, heterozygous mice also showed higher synovitis scores than wild-type mice (MD = 1.792, 95% CI: 0.2670–3.316; P = 0.0182). Interestingly, genotype-related differences were also detected in non-operated contralateral joints, with higher synovitis scores in *Agc1^CreERT2/CreERT2^* compared with *Agc1^+/+^*mice at 4- and 8-w post-DMM with an MD of 1.315 (95% CI: [0.1375, 2.492], *P* = 0.0256) and 2.254 (95% CI: [0.6833, 3.825], *P* = 0.0034). A similar difference was observed between the *Agc1^CreERT2/+^* and *Agc1^CreERT2/CreERT2^*knee joints in the 8-w post-DMM group, with an MD of 1.601 (95% CI: [0.1401, 3.062], *P* = 0.0291). By 12-w post-DMM, genotype-associated differences in synovitis persisted but no longer reached statistical significance [Fig. 4(B)].

**Figure 4.**
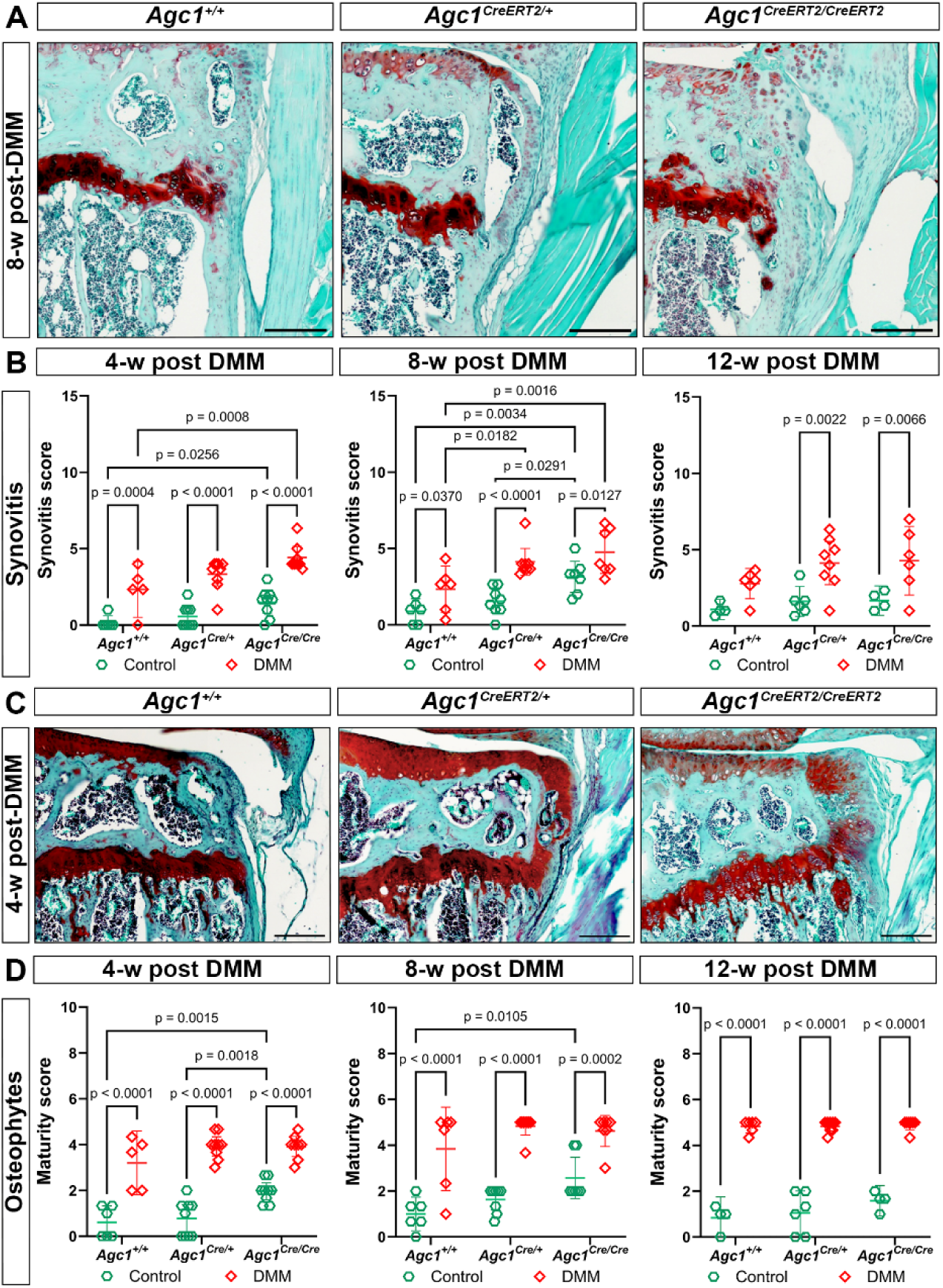
Histological evaluation of synovitis and osteophyte maturation after DMM surgery. (A) Representative Safranin O/Fast Green stained knee joint sections showing synovial changes at 8-w post-DMM in *Agc1^+/+^*, *Agc1^CreERT2/+^*, and *Agc1^CreERT2/CreERT2^* mice. (B) Synovitis scores at 4-, 8-, and 12-w post-DMM; the contralateral knee served as the control. (C) Representative sections illustrating osteophyte formation at 4-w post-DMM across the indicated genotypes. (D) Modified osteophyte maturity scores at 4-, 8-, and 12-w post-DMM; contralateral knees served as controls. The graph data are shown as mean ± 95% CI. Statistical comparisons were performed using two-way ANOVA followed by Tukey’s multiple-comparisons post hoc test (n ≥ 4). Exact p values are shown in the plots. Scale bar = 200 μm.

Osteophyte maturation was influenced predominantly by the DMM-rather than the genotype [Fig. 4(C, D)]. Osteophyte maturity scores [Fig. 4(D)] were significantly higher in DMM-operated joints than contralateral controls across all genotypes and time points. However, differences were also observed among non-operated contralateral joints, with higher scores in homozygous than in wild-type mice at 4- and 8-w (MD = 1.352, 95% CI: 0.4808–2.223, P = 0.0015; and MD = 1.571, 95% CI: 0.3285–2.814, P = 0.0105, respectively). Additionally, homozygous mice displayed higher osteophyte maturity scores than heterozygous mice at 4-w post-DMM (MD = 1.185, 95% CI: 0.4061–1.964; P = 0.0018).

### *Agc1^CreERT2/CreERT2^* mice display pronounced alterations in the subchondral bone structures of the knee

Quantitative assessment of trabecular architecture in the medial subchondral bone at 8-w post-DMM is shown in Fig. 5(A). Trabecular thickness (Tb.Th) was modestly reduced in *Agc1^CreERT2/CreERT2^* mice compared with wild-type (MD = −21.91%, 95% CI: [-43.71, −0.1098], *P* = 0.0486). Trabecular separation (Tb.Sp) slightly increased following DMM in all genotypes but reached statistical significance only in *Agc1^+/+^* mice (MD = 29.94%, 95% CI: [2.303, 57.58], *P* = 0.0343).

**Figure 5.**
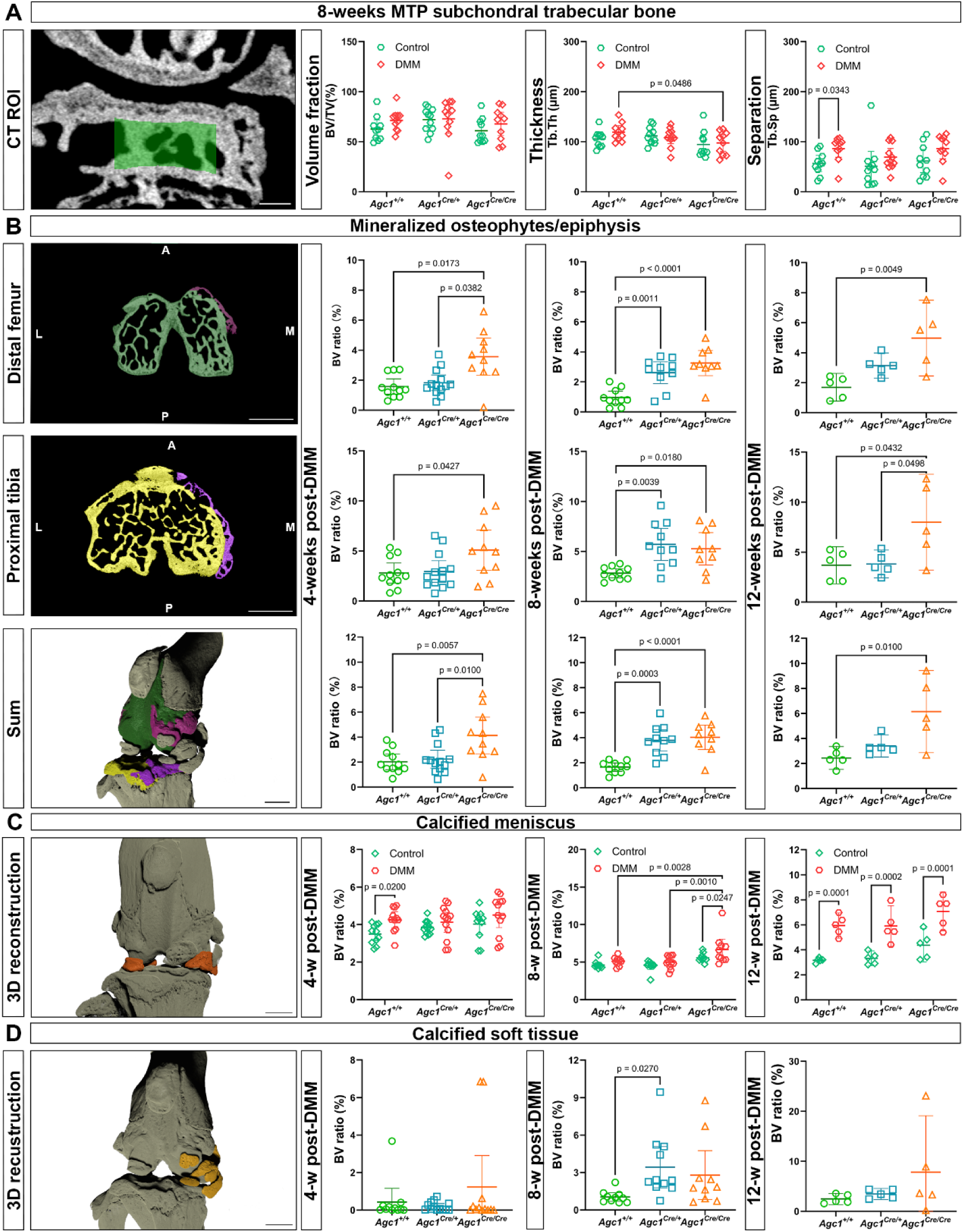
Micro-CT analysis of the knee joint. (A) Representative µCT image of the medial compartment of the knee joint 8-w post-DMM showing the volume of interest (green rectangle) at the subchondral bone trabecular area of the medial tibial plateau and quantitative analysis of the bone volume to total volume (BV/TV), trabecular thickness (Tb.Th) and trabecular separation (Tb.Sp) in *Agc1^+/+^*, *Agc1^CreERT2/+^* and *Agc1^CreERT2/CreERT2^* operated (DMM) and non-operated (control) mice. Scale bar = 200 µm. The graph data are shown as mean ± 95% CI; statistical comparisons among genotypes were performed using two-way ANOVA followed by Tukey’s multiple-comparisons post hoc test (n ≥ 10). (B) Representative 2D and 3D µCT images of the knee joint displaying the osteophytes in purple and the epiphyseal bone in green and yellow. Abbreviations: A, anterior; P, posterior; M, medial; L, lateral. Bar = 1 mm. Quantitative analysis of BV ratio of mineralized osteophytes/attached epiphysis at 4-, 8- and 12-w post-DMM in *Agc1^+/+^*, *Agc1^CreERT2/+^*and *Agc1^CreERT2/CreERT2^* mice. (C) Representative three-dimensional µCT images of calcified menisci post-DMM and quantitative analysis of BV ratio of calcified meniscus/epiphysis at 4-, 8- and 12-w post-DMM and non-operated knee joint in *Agc1^+/+^*, *Agc1^CreERT2/+^*and *Agc1^CreERT2/CreERT2^* mice. The graph data are shown as mean ± 95% CI; statistical comparisons among genotypes were performed using two-way ANOVA followed by Tukey’s multiple-comparisons post hoc test (n ≥ 5). (D) Representative three-dimensional µCT images of abnormal calcified soft tissue and quantitative analysis of BV ratio of abnormal calcified soft tissue/epiphysis at 4-, 8-w and 12-w post-DMM knee joint in *Agc1^+/+^*, *Agc1^CreERT2/+^* and *Agc1^CreERT2/CreERT2^* mice. Scale bar = 1 mm. Data are shown as mean ± 95% CI. One-way ANOVA with Tukey’s post hoc test was used unless otherwise indicated; the distal femur dataset at 4-w post-DMM (Fig. 5B) and the 8-w post-DMM dataset (Fig. 5D) were analyzed using Welch’s one-way ANOVA with Dunnett’s T3 multiple comparisons (n ≥ 5). Exact p values are shown where indicated.

Next, we quantified ectopic calcifications, including osteophytes, meniscal calcification, and calcified periarticular soft tissues. Osteophytes [Fig. 5(B)] developed in all genotypes and at all time points, but were more pronounced in *Agc1^CreERT2/CreERT2^*mice. At 4-w post-DMM, *Agc1^CreERT2/CreERT2^* mice had a significantly higher bone volume (BV) ratio in the femur than *Agc1^+/+^*and *Agc1^CreERT2^*^/*+*^ mice, with MDs of 1.982% (95% CI [0.3469, 3.617], *P* = 0.0173) and 1.734% (95% CI [0.08934, 3.380], *P* = 0.0382), respectively. In the tibia, *Agc1^CreERT2/CreERT2^*mice also had a higher value compared to wild type, with an MD of 2.286%, 95% CI [0.06454, 4.508], *P* = 0.0427. In the sum analysis, *Agc1^CreERT2/CreERT2^*mice showed higher BV ratio compared to both *Agc1^+/+^* and *Agc1^CreERT2/+^*mice with MDs of 2.105% (95% CI [0.5646, 3.644], *P* = 0.0057) and 1.893% (95% CI [0.4109, 3.376], *P* = 0.0100), respectively. At 8-w post-DMM, both *Agc1^CreERT2/CreERT2^* and *Agc1^CreERT2/+^*mice had higher BV ratios than *Agc1^+/+^* mice. In *Agc1^CreERT2/CreERT2^*mice, the MDs were 2.300% in the femur (95% CI [1.255, 3.345], *P* < 0.0001), 2.435% in the tibia (95% CI [0.3779, 4.492], *P* = 0.0180), and 2.381% in the combined analysis (95% CI [1.198, 3.563], *P* < 0.0001). The corresponding MDs for *Agc1^CreERT2/+^* mice were 1.659% in the femur (95% CI [0.6423, 2.677], *P* = 0.0011), 2.878% in the tibia (95% CI [0.887753, 4.880], *P* = 0.0039), and 2.137% in the sum (95% CI [0.9863, 3.288], *P* = 0.0003). By 12-w, the values of the homozygous mice reached their peak and were significantly elevated compared with wild-type mice in the femoral compartment, with an MD of 3.280% (95% CI [1.074, 5.486], *P* = 0.0049); relative to both wild-type and heterozygous mice in the tibial compartment with MDs of 4.304% (95% CI [0.1294, 8.478], *P* = 0.0432), and 4.177% (95% CI [0.003129, 8.352], *P* = 0.0498), respectively; and compared with wild-type mice in the combined analysis with an MD of 3.708% (95% CI [0.9376, 6.479], *P* = 0.0100).

Meniscal and periarticular soft tissue mineralization was localized predominantly to the medial compartment [Fig. 5(C, D)]. At 8-w post-DMM, meniscal mineralization was significantly greater in DMM-operated *Agc1^CreERT2/CreERT2^*mice compared with both *Agc1^+/+^* and *Agc1^CreERT2/+^*mice, with MDs of 1.556% (95% CI [0.4790, 2.633], *P* = 0.0028) and 1.669% (95% CI [0.6172, 2.721], *P* = 0.0010). In addition, homozygous mice exhibited a significant difference between DMM-operated and contralateral joints, with an MD of 1.061% (95% CI [0.1406, 1.981], *P* = 0.0247). At 12-w, the BV ratio of calcified menisci in DMM-operated *Agc1^CreERT2/CreERT2^*mice remained numerically higher than those in DMM-operated *Agc1^+/+^*and *Agc1^CreERT2/+^* mice, with MDs of 1.126% (95% CI [-0.3654, 2.618]) and 1.164% (95% CI [−0.3272, 2.656]); however, neither difference reach statistical significance. On the other hand, at this late stage, robust increases in meniscal calcification were observed in DMM-operated mice compared with contralateral joints in all genotypes.

With respect to periarticular soft tissues, µCT analysis revealed no overt ectopic calcification at 4-w post-DMM, apart from occasional isolated outliers [Fig. 5(D)]. At subsequent time points, *Agc1^CreERT2/+^* (8-w) and *Agc1^CreERT2/CreERT2^*(8- and 12-w) mice demonstrated increased BV ratios of calcified soft tissues compared with wild-type mice. However, the only significant post-DMM genotype comparison occurred between *Agc1^+/+^* and *Agc1^CreERT2/+^*mice at 8-w, with an MD of −2.383% (95% CI [-4.497, −0.2681], *P* = 0.0270).

### *Agc1^CreERT2/CreERT2^* mice exhibit elevated cartilage catabolism

Next, we evaluated the expression of key cartilage-degrading proteinases, including MMP-3, MMP-9, MMP-13, ADAMTS-4, and ADAMTS-5, together with degradation neoepitopes of type II collagen (C1,2C) and aggrecan (VDIPEN generated by MMPs, NITEGE generated by aggrecanase activity), as well as markers of chondrocyte secretory activity (type II collagen and lubricin). Among these, C1,2C, NITEGE, MMP-13 and ADAMTS-5 displayed genotype-dependent differences [Fig. 6 (A, B)]. At 4-w post-DMM, levels of NITEGE were increased in *Agc1^CreERT2/+^*and *Agc1^CreERT2/CreERT2^* mice, and ADAMTS-5 expression was elevated in *Agc1^CreERT2/CreERT2^* mice [Fig. 6 (A)]. At 8-w post-DMM, C1,2C, NITEGE, MMP-13 and ADAMTS-5 immunostaining was increased in *Agc1^CreERT2/CreERT2^*mice compared to *Agc1^+/+^* and *Agc1^CreERT2/+^* mice [Fig. 6(B)].

**Figure 6.**
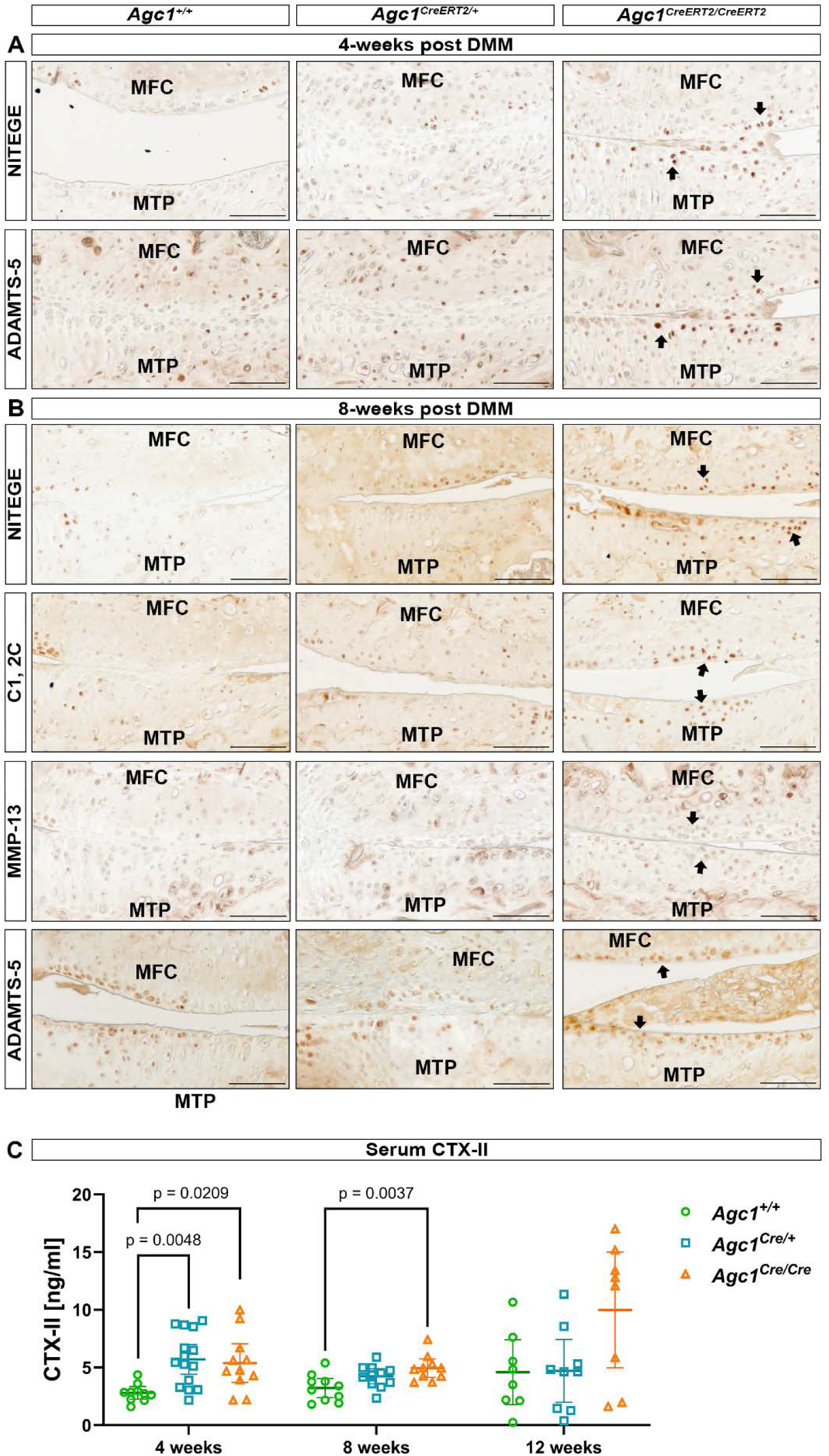
Altered expression of cartilage catabolic markers and serum CTX-II levels post-DMM. (A,B) Representative immunohistochemical staining in the MFC and MTP of AC of *Agc1^+/+^*, *Agc1^CreERT2/+^*, and *Agc1^CreERT2/CreERT2^*. (A) NITEGE and ADAMTS-5 at 4-w post-DMM. (B) NITEGE, C1,2C, MMP-13, and ADAMTS-5 at 8-w post-DMM. Arrows indicate areas of prominent immunoreactivity and cartilage lesion. Scale bars = 50 µm. (C) Serum CTX-II concentrations measured by ELISA at 4-, 8-, and 12-w post-DMM. Individual data points and mean ± 95% CI are shown. Comparisons between genotypes were performed using Welch’s ANOVA followed by Dunnett’s T3 multiple-comparisons test at 4 and 12 weeks, and ordinary one-way ANOVA followed by Tukey’s multiple-comparisons test at 8 weeks (n ≥ 8). Exact *P* values are indicated in the graph.

Serum concentrations of CTX-II, a biomarker of type II collagen degradation, were measured by ELISA at 4-, 8-, and 12-w post-DMM. At all time points, *Agc1^CreERT2/CreERT2^* mice displayed the highest CTX-II levels [Fig. 6(C)]. At 4-w post-DMM, CTX-II concentrations were significantly higher in *Agc1^CreERT2/CreERT2^*than *Agc1^+/+^* mice, with an MD of 2.580 ng/ml (95% CI [0.3433, 4.816], *P* = 0.0209). The difference also persisted at 8-w post-DMM, with an MD of 1.723 ng/ml (95% CI: [0.5266, 2.918], *P* = 0.0037). Notably, heterozygous mice also showed increased CTX-II levels relative to wild-type mice at 4-w, with an MD of 2.906 ng/ml (95% CI: [0.8163, 4.996], *P* = 0.0048). At 12-w, CTX-II remained numerically highest in homozygous mice but did not differ significantly from wild-type or heterozygous mice.

### Chondrocyte apoptosis is increased in *Agc1^CreERT2/CreERT2^* mice during PT-OA

TUNEL staining at 4- and 8-w post-DMM revealed increased apoptotic cell numbers in *Agc1^CreERT2/CreERT2^* mice compared to the other genotypes [Fig. 7(A)]. Quantification at 4-w post-DMM demonstrated significantly increased apoptosis in the uncalcified zone of the AC in *Agc1^CreERT2/CreERT2^*mice compared to *Agc1^+/+^* and *Agc1^CreERT2/+^* littermates, with MDs of 3.183% (95% CI [0.8221, 5.545], *P* = 0.0080) and 3.365% (95% CI [1.156, 5.574], *P* = 0.0031), respectively. When combining the uncalcified and calcified zones, *Agc1^CreERT2/CreERT2^* mice exhibited a significantly higher proportion of TUNEL-positive cells compared with wild-type and heterozygous mice (MD = 3.098%, 95% CI: 0.7230– 5.473, P = 0.0101; and MD = 3.196%, 95% CI: 0.9749–5.418, P = 0.0049, respectively). At 8-w post-DMM, apoptotic rates in *Agc1^CreERT2/+^* mice approached those observed in *Agc1^CreERT2/CreERT2^* mice, with comparable values in the uncalcified zone, calcified zone, and combined analysis. Although one-way ANOVA identified a significant overall difference in the combined apoptotic ratio among genotypes (F = 3.683, P = 0.0483, R^2^ = 0.3152), post hoc comparisons did not reveal significant differences between individual groups. Significant pairwise differences between genotypes were detected only in the uncalcified zone, where both heterozygous and homozygous mice exhibited higher TUNEL-positive cell ratios than wild-type mice. (MD = 3.090%, 95% CI [0.6592, 5.521], P = 0.0124; and MD = 3.571%, 95% CI [1.049, 6.094], P = 0.0057, respectively) [Fig. 7(B)].

**Figure 7.**
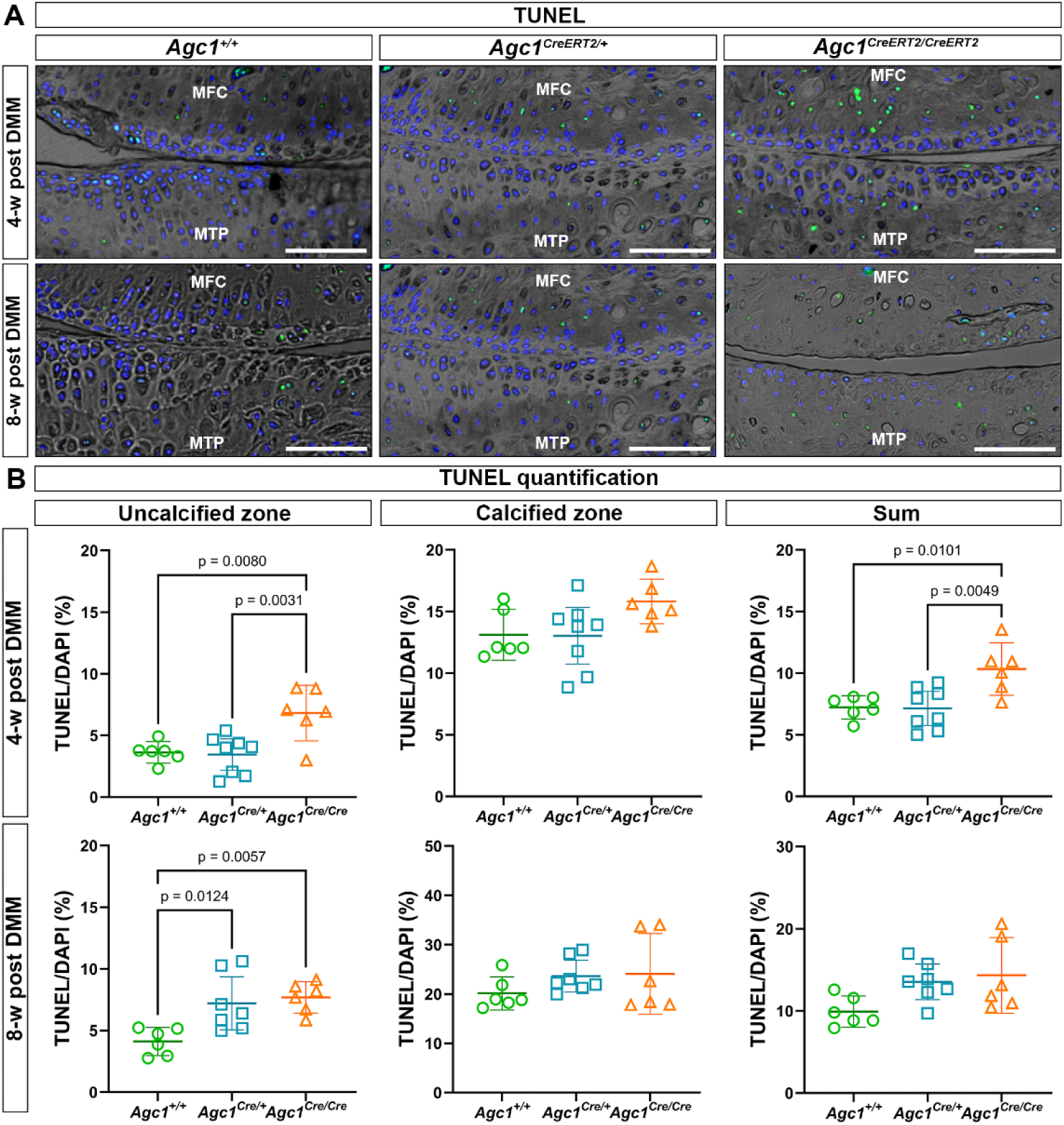
Chondrocyte apoptosis assessed by TUNEL assay in *Agc1^+/+^*, *Agc1^CreERT2/+^*and *Agc1^CreERT2/CreERT2^* mice at 4- and 8-w post-DMM. (A) Representative merged images of phase contrast and fluorescent TUNEL (green) and DAPI (blue) staining of the medial compartment of the knee joint. Scale bar = 100 µm. (B) Quantification of apoptosis rate (TUNEL-positive cells/DAPI positive cells) at 4- and 8-w post-DMM in *Agc1^+/+^*, *Agc1^CreERT2/+^*and *Agc1^CreERT2/CreERT2^* mice. Data are performed as means ± 95% CI; and comparisons of genotypes were conducted using one-way ANOVA with Tukey’s post hoc test, except for calcified zone at 8-w when Welch’s one-way ANOVA with Dunnett’s T3 multiple comparisons were used (n ≥ 6). Exact p values are shown where indicated.

### DMM leads to an increase in the stiffness of AC zones in all genotypes

Because homozygous mice exhibited altered AC biomechanics at baseline, we next investigated whether these differences persisted following DMM surgery. Overall, Young’s modulus increased continuously from the superficial to the deep cartilage zone in both DMM-operated and contralateral knees [Fig. 8(A,B)]. Across all zones, *Agc1^CreERT2/CreERT2^* cartilage remained stiffer than wild-type cartilage under both conditions. Compared to the non-operated control group, the deep zone exhibited the most considerable change in stiffness after DMM. In wild type, E1 increased by 177% and E2 by 132% in the deep zone after DMM compared with the contralateral joint. This trend was less pronounced in *Agc1^CreERT2/+^* and *Agc1^CreERT2/CreERT2^*mice, yet the increase remained substantial. In the middle zone of wild-type, DMM increased E1 and E2 by 80% and 53%, respectively. *Agc1^CreERT2/+^*mice showed a similar pattern, with E1 increasing by 29% and E2 by 60%, whereas *Agc1^CreERT2/CreERT2^* mice displayed only a slight increase. Interestingly, in the superficial zone, DMM had a greater impact on heterozygous mice, with E1 increasing by 67%, E2 by 116%, and the range of Young’s modulus values increased by 62% (0.01-5.85 MPa vs. 0.00-3.61 MPa) compared to contralateral cartilage. These results indicate that DMM induces an increase in AC stiffness across all cartilage zones and genotypes, with both *Agc1^CreERT2/+^*and *Agc1^CreERT2/CreERT2^* mice showing higher stiffness in the superficial and middle zones than wild-type mice, and *Agc1^CreERT2/CreERT2^*mice exhibiting the greatest stiffness in all zones compared with other genotypes.

**Figure 8.**
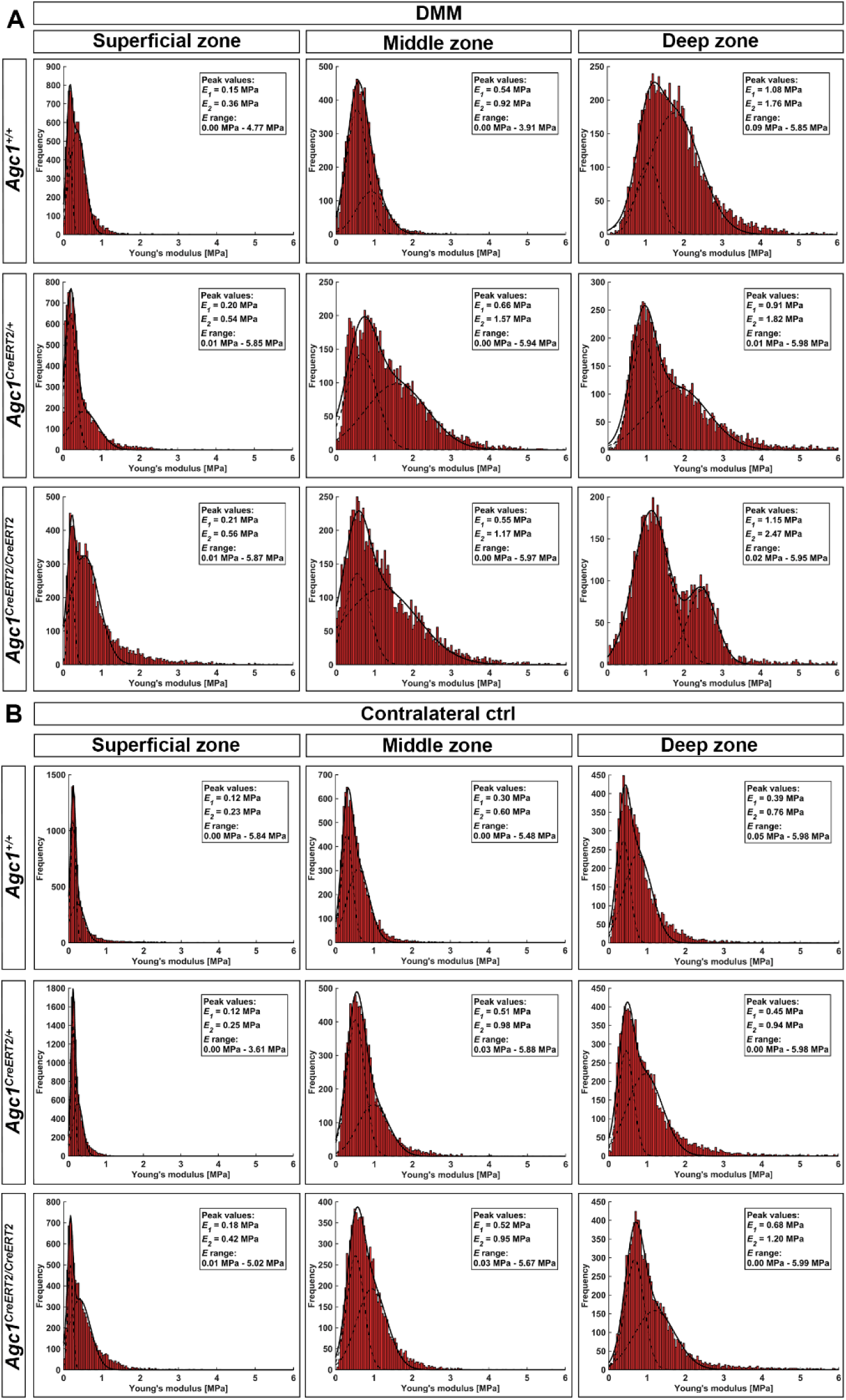
IT-AFM measurements of the superficial, middle and deep zones of the medial tibial plateau of DMM operated (A) and contralateral, non-operated (B) knee joints 4-w post-DMM. In each histogram, the solid line represents the combination of two Gaussian functions, while the dashed lines indicate individual fits. The first and second stiffness peaks (E1 and E2) correspond to the proteoglycan gel and the collagen fibrils, respectively. n = 3 animals per genotype.

## Discussion

Aggrecan, the major sulfated proteoglycan of ECM of the growth plate and AC, is essential for the proper structure and biomechanical function of hyaline cartilage. Structural and functional mutations in the gene encoding aggrecan cause skeletal dysplasia with or without joint abnormalities, and reduced expression of the aggrecan protein is a critical determinant (20) and hallmark (21) of the onset and progression of OA. In humans, heterozygous nonsense or frameshift *ACAN* variants cause spondyloepiphyseal dysplasia Kimberley type, characterized by short stature and severe early-onset OA (6,9), whereas missense variants underlie familial osteochondritis dissecans with impaired aggrecan secretion or ECM interactions (7,22). A study of 20 families with heterozygous *ACAN* mutations and autosomal dominant short stature identified 12 families with early-onset OA, predominantly in the knee joint (23). While the adult short stature phenotype is likely due to haploinsuffiency of aggrecan, the susceptibility to early-onset OA might be modulated by additional genetic or environmental factors. In animals, homozygous loss-of-function aggrecan mutations cause lethal chondrodysplasia in mice (cartilage matrix deficiency, *cmd*), chicken and cattle (24–26), whereas heterozygous *cmd/+* mice develop postnatal proportional dwarfism and spinal disc degeneration (27). We and others have described that homozygous *Agc1^CreERT2/CreERT2^*mice develop dwarfisms by 1 month (11) and severe knee OA by 1 year (12). Here, using the injury-induced DMM model, we demonstrate that reduced sGAG content in *Agc1^CreERT2/CreERT2^*mice increases susceptibility of the entire knee joint to trauma-induced OA, resulting in more severe cartilage degeneration, synovitis, soft tissue pathology and bone changes.

Mouse models of human aggrecanopathies exhibit varying degrees of aggrecan deficiency, resulting in a spectrum of skeletal phenotypes. In *cmd* mice, aggrecan mRNA in embryonic limb cartilage is reduced to 81% in *cmd/+* and 41% in *cmd/cmd* mice (27), whereas single-cell RNA sequencing reported an approximately 50% reduction in *Acan* mRNA in *cmd/+* mice (28,29). Chondroitin sulfate in spinal cartilage in 3-month-old *cmd/+* mice was reduced to 87% of the wild-type (27). We recently generated the aggrecan insertion mutant *Acan^iE5^*mouse, in which a *loxP* insertion in exon 5 causes aggrecan deficiency similar to that in *cmd* mice (30). *Acan^iE5/iE5^*mice die at birth with severe chondrodysplasia, complete loss of aggrecan core protein and cartilage sGAG levels reduced to 21.59% of wild-type, whereas heterozygous *Acan^iE5/+^* mice show normal skeletal development despite a reduction in cartilage sGAG content to 86.35%. In contrast, *cmd/+* mice show growth failure as early as 3-4 weeks after birth (27,28) and age-associated intervertebral disc degeneration at 1 year (27), but neither heterozygous model develops spontaneous OA or has been examined in post-traumatic OA models. In the present study, *Agc1^CreERT2/CreERT2^*mice showed a reduction in sGAG content to 75% in E18.5 epiphyseal cartilage and reduced aggrecan immunostaining in 3-month-old AC. Consistent with this, Rashid et al. reported a 50% reduction in aggrecan protein and a 47% decrease in proteoglycan content in the knee cartilage of 3-month-old *Agc1^CreERT2/CreERT2^*mice (11). These changes are associated with dwarfism from 1 month of age (11,12), a 27% reduction in AC thickness at 3 months, and severe cartilage degeneration by 12 months (12). Importantly, before DMM surgery, 3-month-old *Agc1^CreERT2/CreERT2^*animals showed no evidence of spontaneous AC degeneration, although synovitis and osteophyte maturation scores tended to be higher in both *Agc1^CreERT2/CreERT2^*and *Agc1^CreERT2/+^* mice than in wild-type controls.

In the present study, reduced aggrecan levels in *Agc1^CreERT2/CreERT2^*mice accelerated DMM-induced cartilage degradation, with significantly higher OARSI scores in the MTP at 4-w post-DMM. By 12-w, both the *Agc1^CreERT2/CreERT2^*and *Agc1^CreERT2/+^* mice exhibited more severe cartilage degeneration than wild-type mice. The integrity of the aggrecan-hyaluronan aggregate and collagen fibril network is essential for biomechanical function of the AC (12,31,32). Although proteoglycan loss and collagen degradation are central features of OA, their temporal sequence depends on the initiating insult. Physiological loading of cartilage with focal defects induces early proteoglycan depletion followed by collagen degradation, whereas injurious loading of intact cartilage or physiological loading of degenerated cartilage initially damages the collagen network, accelerating subsequent proteoglycan loss (33). Likewise, cyclic overloading of healthy cartilage causes proteoglycan loss without collagen damage, whereas collagen-compromised cartilage exhibits markedly greater proteoglycan depletion under mechanical overload (34). In IL-1-stimulated bovine cartilage explants, proteoglycan depletion also precedes collagen degradation, and inhibition of aggrecanases prevents subsequent MMP-mediated collagen breakdown, supporting a protective role for aggrecan against collagen proteolysis (35). Consistent with this concept, DMM increased serum CTX-II levels in both *Agc1^CreERT2/CreERT2^*and *Agc1^CreERT2/+^* mice, as well as immunoreactivity of C1,2C, ADAMTS-5 and MMP-13 in the AC of *Agc1^CreERT2/CreERT2^* mice compared with wild-type, implicating that the protective role of aggrecan against collagen breakdown is impaired in hypomorphic mice. Furthermore, increased ADAMTS-5 expression and accumulation of its cleavage neoepitope NITEGE at 8-w post-DMM, suggest that suboptimal aggrecan levels make AC vulnerable to aggrecanase activation and matrix degradation. As ADAMTS-5 is a major aggrecanase in OA, and its genetic deletion attenuates cartilage destruction in the DMM model (36), these findings indicate that reduced aggrecan not only compromises cartilage biomechanics but also enhances susceptibility to catabolic matrix remodeling following joint injury.

During early OA, aggrecan content initially increases but subsequently declines due to enhanced proteolysis, contributing to progressive cartilage degeneration (37,38). Aggrecan-link protein-hyaluronan aggregates generate the cartilage swelling pressure that is balanced by the collagen fibrillar network. Their retention within the ECM depends on interactions with adaptor proteins, including decorin, biglycan and matrilins, and disruption of these interactions compromises matrix integrity and increases OA susceptibility. Accordingly, decorin deficiency reduces aggrecan and sGAG content, alters collagen nanostructure (39), and accelerates DMM-induced OA (40). Likewise, matrilin-1 deficiency decreases aggrecan and collagen expression and exacerbates OA after DMM (41). In humans, the matrilin-3 T303M mutation is associated with hand OA (42), while the corresponding murine T298M mutation increases proteoglycan extractability and PT-OA severity, suggesting that impaired anchorage of aggrecan within the collagen network increases cartilage vulnerability to OA (43). These studies reinforce the concept that maintenance of an intact aggrecan network is critical for preserving AC function and limiting both age-related and PT-OA.

We previously reported that aggrecan insufficiency increases the ECM stiffness across all zones of AC in 6-month-old *Agc1^CreERT2/CreERT2^*mice (12). Here, increased superficial zone nano-stiffness was already evident in 3-month-old homozygous mice. Consistent with previous nano-indentation studies showing increased stiffness of collagen- and proteoglycan-rich matrix 2-w after DMM, with stiffness increasing from the superficial to the deep zone (44), we observed a similar depth-dependent increase in nano-stiffness and higher values in DMM than control cartilage across all genotypes at 4-w post-DMM. In contrast, micro-indentation detected cartilage softening after DMM (3), suggesting that proteoglycan loss may differentially affect cartilage biomechanics depending on indentation scale. Similarly, cathepsin D-mediated proteoglycan depletion increased nano-stiffness in porcine cartilage (45), and aggrecan-deficient *Acan^iE5/iE5^* mice exhibited marked stiffening of the cartilaginous endplate (30). Together, these findings demonstrate that reduced aggrecan content results in increased nano-stiffness in the superficial and middle zone of both *Agc1^CreERT2/CreERT2^*and *Agc1^CreERT2/+^* mice compared with wild-type mice during PT-OA, highlighting the importance of aggrecan in maintaining cartilage biomechanical properties.

Increased ECM stiffness may enhance mechanical stress on chondrocytes, thereby promoting apoptosis and accelerating cartilage degeneration (46). Consistent with this, deletion of the aggrecan coding sequence in *Acan^cmd-Bc^* mice causes perinatal lethal chondrodysplasia accompanied by increased chondrocyte apoptosis in developing limb cartilage, whereas transgenic restoration of aggrecan largely rescues both the skeletal phenotype and cell survival (47). We found significantly increased chondrocyte apoptosis in the non-calcified AC of *Agc1^CreERT2/CreERT2^* mice at 4- and 8-w post-DMM, and in *Agc1^CreERT2/+^* mice at 8-w, but was unchanged in non-operated 3-month-old mice. This suggests that partial aggrecan deficiency alone is insufficient to induce apoptosis, whereas mechanical injury exacerbates biomechanical and catabolic changes, increasing chondrocyte vulnerability. The accelerated proteoglycan loss and collagen degradation observed during DMM, particularly in *Agc1^CreERT2/CreERT2^* mice, are likely to contribute to this response, as disruption of both the aggrecan and collagen II networks promotes chondrocyte apoptosis (48–50), underlining the importance of an intact ECM in maintaining chondrocyte survival.

We observed pronounced exacerbation of synovitis, osteophyte formation, meniscal and periarticular soft tissue ossification in *Agc1^CreERT2/CreERT2^*mice. Acute joint injury rapidly induces synovial inflammation and tissue remodeling. DMM-induced joint instability could also promote pathological changes in the meniscus, ligaments and synovial capsule that may progress through chondrogenesis, calcification and bone formation (51–53). Consistent with previous studies showing that synovitis precedes cartilage degeneration after DMM and subsequently correlates with cartilage destruction and aggrecan loss (54), *Agc1^CreERT2/CreERT2^* mice displayed increased synovitis before surgery and significantly more severe synovitis at 4- and 8-w post-DMM, while *Agc1^CreERT2/+^* mice developed significantly increased synovitis at 8-w. Although histological osteophyte maturation scores were comparable between genotypes, homozygous hypomorphic mice developed significantly more mineralized osteophytes at all post-DMM time points. Recent studies demonstrated that synovial mesenchymal progenitor cells (MPCs) express aggrecan, and that intra-articular aggrecan administration attenuates cartilage degeneration in a PT-OA model and increases cartilage repair in a focal injury model (55). Furthermore, transplantation of synovial MPCs isolated from wild-type mice enhanced cartilage regeneration more effectively than MPCs from *Agc1^CreERT2/CreERT2^*mice (55). These findings support a broader protective role for synovial-derived aggrecan in limiting post-traumatic joint pathology, suggesting that reduced aggrecan deposition may exacerbate synovial inflammation, ectopic mineralization and OA progression. However, synovitis is rarely recognized in aggrecanopathies caused by *ACAN* mutations associated with OCD, short stature and early-onset osteoarthritis, with only a single case describing synovial thickening and elbow inflammation in a child carrying an *ACAN* missense mutation (56).

In conclusion, hypomorphic *Agc1^CreERT2/CreERT2^* mice (14) exhibit reduced sGAG and aggrecan protein levels in cartilaginous tissues, resulting in exacerbated PT-OA following DMM. Reduced aggrecan compromises the biomechanical properties of articular cartilage and promotes more severe synovitis, cartilage degeneration, chondrocyte apoptosis, and ectopic calcification after joint injury. Importantly, these pathological changes were not restricted to homozygous mice. Heterozygous *Agc1^CreERT2/+^*mice also developed significantly increased synovitis, osteophyte and soft tissue mineralization, and AC degeneration at 8- and 12-w post-DMM. *Agc1^CreERT2^*mice are widely used for tamoxifen-inducible, cartilage-specific gene deletion to study gene function in osteoarthritis. Because homozygous *Agc1^CreERT2/CreERT2^*mice develop dwarfism and age-associated OA, it is a standard practice to use the heterozygous *Agc1^CreERT2/+^* mice for generation of conditional knockouts (11,12). Our findings demonstrate that both homo- and heterozygous *Agc1^CreERT2^* mice display advanced joint pathology following DMM, supporting a critical role for aggrecan dosage in determining susceptibility to PT-OA. Therefore, studies using this *Cre* line to investigate gene function in PT-OA should carefully consider the contribution of the *Agc1^CreERT2^* allele itself when interpreting disease severity.

## Author contribution

Attila Aszodi, Paolo Alberton and Hauke Clausen-Schaumann developed study concept and design, analyzed and interpreted data, revised the draft paper and approved the final version. Xujia Wang performed experiments, analyzed data, generated figures and wrote the paper. Bastian Hartmann performed AFM experiments, analyzed and interpreted data and contributed to writing the manuscript. Reinhild Hofmann, Shenxi Zhong, Xiangqing Meng, Heidrun Grondinger and Zsuzsanna Farkas performed experiments. All authors contributed to the article and approved the submitted version.

